# Revisiting graph-based approaches for small protein analysis: Insights from anti-CRISPR protein networks

**DOI:** 10.1101/2025.10.31.685784

**Authors:** Michelle Ramsahoye, Mirela Alistar

**Affiliations:** Department of Computer Science, University of Colorado Boulder, Boulder, Colorado, USA; ATLAS Institute, University of Colorado Boulder, Boulder, Colorado, USA

## Abstract

Bacteriophage anti-CRISPR (Acr) proteins have the potential to reduce off-target effects of genome editing by inactivating the CRISPR-Cas bacterial defense. The current challenge lays in their functional annotation, as Acr proteins have high structural diversity and low sequence similarity, thus rendering common homology-based methods unfit. Recent solutions use deep learning models such as graph convolutional networks that take protein networks as the data input. In an effort to understand whether these new solutions are fit for niche, sparsely annotated proteins, we focus on 3 Acr proteins (AcrIF1, AcrIIA1, and AcrVIA1) as a case study. For each, we create protein contact networks (PCNs) and residue interaction graphs (RIGs) based on existing network theory and methodology. We characterize and analyze these protein networks by comparing how each network architecture affects values of small-worldliness. We reexamine a previous method that focused on using node degree, closeness centralities, and residue solvent accessibility to predict functional residues within a protein via a Jackknife technique. We discuss the implications of the construction of these networks based on how the structure information is acquired. We demonstrate that functional residues within small proteins cannot be reliably predicted with the Jackknife technique, even when provided with a curated dataset containing representative standardized values for degree and closeness centrality. We show that functional residues within these small proteins have low degrees within both PCNs and RIGs, thus making them susceptible to the known degree bias towards high degree nodes present in using graph convolutional networks. We discuss how understanding the data can be used to further improve deep learning approaches for small proteins.

**Author summary:** A bacteria’s CRISPR-Cas defense system acts as security guard against viruses like bacteriophages. By storing pieces of viral DNA as records, it can recognize and defend the bacteria against threats. Scientists have adapted this effective record keeping process to perform targeted genome editing. Some bacteriophages have genes that encode for anti-CRISPR (Acr) proteins. The proteins act as a criminal accomplice to the viral DNA, sneaking them in past the bacteria’s security in a variety of ways. There has been increased interest in using these Acr proteins to limit unintended or off-target effects of targeted genome editing. However, Acr proteins are difficult to identify. We changed parts of a previous method that used graph representations of protein structure to determine important amino acids that help that protein perform its function. We applied these methods to three Acr proteins to determine whether we observed similar patterns in these graphs. We explain how features of these graph representations of protein structures can affect graph neural networks that use them as input to learn more about proteins.

## Introduction

Anti-CRISPR (Acr) proteins are a unique class of proteins capable of inactivating the CRISPR-Cas bacterial defense system. First discovered in 2013 by the Bondy-Denomy lab within lysogens of *Pseudomonas aeruginosa* PA14 [1], these proteins allow bacteriophages to invade bacteria and introduce their genetic material for replication. Since their discovery, Acr proteins have been proposed as a means to reduce the off-target effects during CRISPR-Cas experiments and thus regulate genome editing [2].

However, identifying new Acr proteins is difficult due to their *high structural diversity* and *low sequence similarity*. Functionally annotating these proteins are an additional challenge as the mechanisms through which they inactivate CRISPR-Cas systems can vary, ranging from DNA mimicry to steric blocking [3]. These challenges are interdependent - commonly used homology-based methods rely on well-annotated reference proteins. When there are very few of these proteins, we limit ourselves to biased discovery of only well-characterized families and any other proteins remain unknown.

In 2015, Graham F. Hatfull described bacteriophages as the “dark matter” of the biosphere due to their high genetic diversity, thus resulting in high functional novelty [4]. Most bacteriophage genes encode for Hypothetical Proteins of Unknown Function (“HPUFs”) [5]. Standard protein function annotations, such as Gene Ontology (GO) terms, are scarce. Of the 543,466 bacteriophage proteins available on the National Center for Biotechnology Information Reference Sequence (NCBI RefSeq) Database (sourced from 5,411 complete genomes via INPHARED [6]), only about 20% of these proteins have GO annotations [7]. Bacteriophage-specific annotations are also lacking. Phamerator [8], which groups protein-coding genes into ’phamilies’ of related sequences via pairwise similarity searches, have been used in curated databases like the Actinobacteriophage Database (also known as PhagesDB). However, even this specialized resource has only 5,586 fully annotated (or ’phamerated’) bacteriophages out of 29,146 entries [9].

Several *in silico* methods exists for protein identification and functional annotation, including homology-based searches [10], large-language models [11], and guilt-by-association approaches [12]. In this work, we focus on network theory-based methods for protein analysis. Network theory uses graphs (mathematical structures of nodes and edges) to model complex relationships. These graph-based approaches been applied to diverse biological macromolecules with examples including RNA motif analysis [13], DNA sequence statistics [14], and protein characterization [15–19]. Here we apply this framework specifically to proteins as shown in Fig. 1, modeling Acr proteins as graphs where **nodes** represent residues and **edges** represent spatial or interatomic interactions.

**Fig 1.**
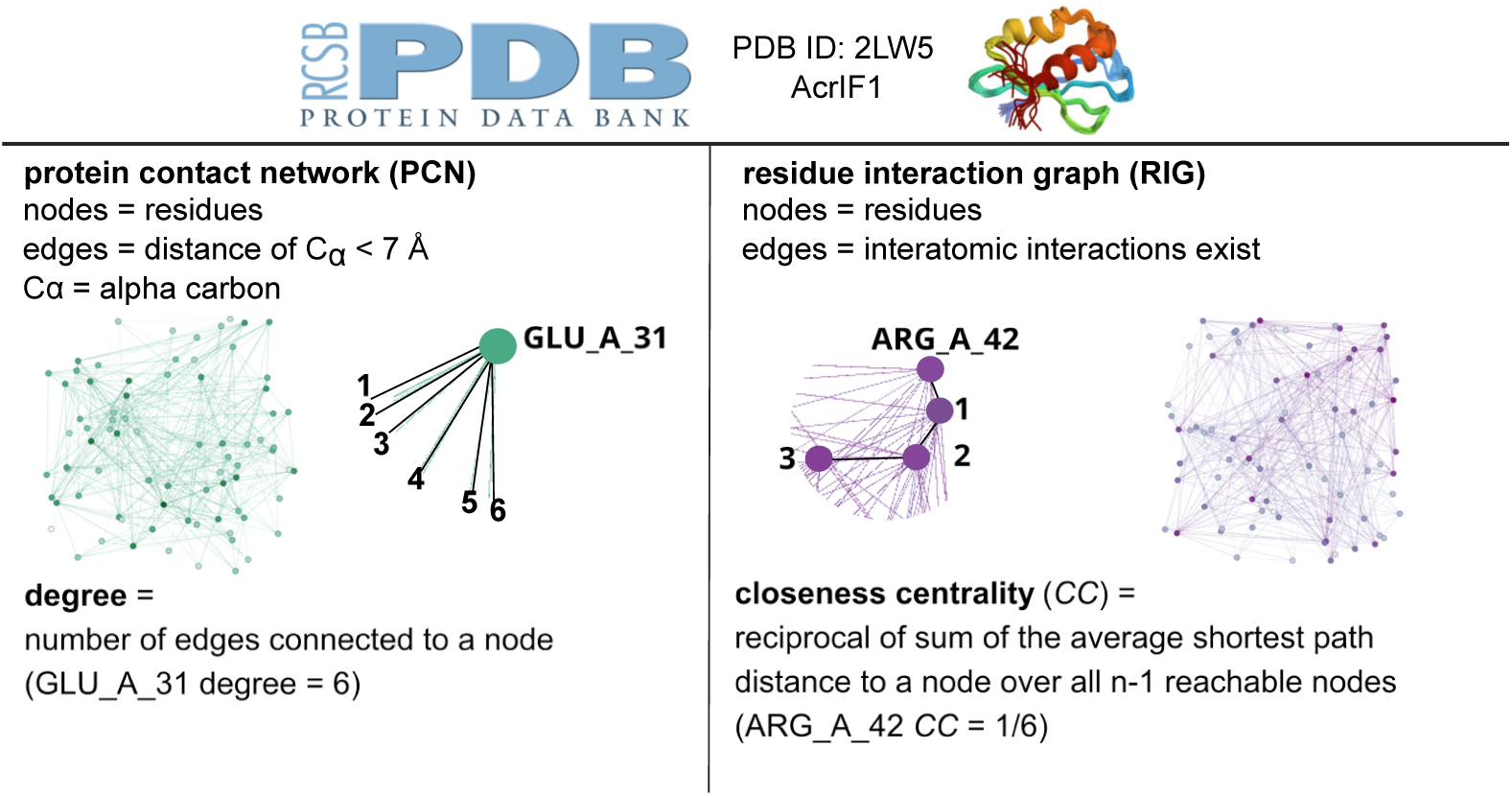
Protein network architectures and definitions. The structure files for all anti-CRISPR proteins are sourced from the Protein Data Bank (PDB). We define a network type of interest (protein contact network/PCN or residue interaction graph/RIG). PCNs define nodes as residues and edges exist where a distance threshold is met between the alpha carbons (*C_α_*) of the residue. An alpha carbon is the central location in the backbone of an amino acid. RIGs define nodes as residues as well, and edges exist where there is an interatomic interaction (hydrogen bonds, Van der Waals forces, etc.) found using PDBe-Arpeggio. Those networks can then be analyzed based on degree and closeness centrality, as previously done in Amitai et al. [21].

The applications of network theory to proteomics has continued into more modern tools such as the Flatiron Institute’s DeepFri model, which uses a graph convolutional network and a large language model to make protein function predictions and assign GO terms [20]. However, like many black box models, the details as to why the model makes its predictions are unknown to the user and likewise, performance of the model is highly dependent on the dataset it is trained on. The training data guides the model – and thus, it can introduce biases, leading to incorrect or unsubstantial conclusions in regards to novel proteins like Acr proteins. Relative to the 200,000 structures found on the Protein Data Bank (PDB) and 254 million protein sequences on the UniProtKB database, Acr protein data is extremely scarce with 122 Acr proteins with 92 different subtypes found to date [3]. This prompts a preliminary question: before deploying models on niche, sparsely annotated proteins, what can we learn about the data itself that might guide future model adaptations?

Application of network theory to protein contact networks (PCNs) and residue interaction graphs (RIGs) can offer mathematical rigor and provide a more transparent means of providing scientific understanding of protein structure and function. Amitai et al. demonstrated that common network analysis methods such as degree and closeness centrality could be used to predict active site residues on individual proteins [21]. This method is largely non-reliant on sequence conservation, comparison to similar structures, or prior knowledge [21]. This method also provides a validated network theory framework for analyzing protein structures, which we adapt here for Acr proteins.

In this work, we apply the network theory framework established by Amitai et al. [21] to three structurally diverse Acr proteins: AcrIF1 (PDB ID: 2LW5), AcrIIA1 (PDB ID: 5Y6A), and AcrVIA1 (PDB ID: 6VRB). While the original methodology focused on monomeric proteins characterized by x-ray diffraction, our work uses monomeric, dimeric, and complex structures derived from solution NMR, x-ray diffraction, and electron microscopy, respectively. As presented in Fig 1, we construct protein contact networks (PCNs) and residue interaction graphs (RIGs) for each protein and characterize them using network metrics including degree, closeness centrality, and residue solvent accessibility. Our analysis reveals that while these Acr protein representations exhibit small-world network properties across three established indices (sigma, omega, and small-world index), certain characteristics deviate from the proposals in the original methodology. We examine the implications of these findings in the context of using structurally diverse and sparsely annotated data as inputs for deep learning models, and discuss how these data properties should inform future model architecture decisions.

## Results

We created network representations for three anti-CRISPR (Acr) proteins: AcrIF1 (PDB ID: 2LW5), AcrIIA1 (PDB ID: 5Y6A), and AcrVIA1 (PDB ID: 6VRB, in complex). The network representations are the protein contact network (PCN) and the residue interaction graph (RIG). More detailed explanations about these representations can be found within the Methods section, but a simplified definition for each is the following:

- Protein contact network (PCN) architecture denotes a node as a residue’s alpha carbon, and the edges exist if two alpha carbons are within 7 angstroms of each other. An alpha carbon is the central location in the backbone of an amino acid.
- Residue interaction graph (RIG) architecture denotes a node as a residue, and edges exist if there are interatomic interactions between the residues (not limited to alpha carbons of the residue). Examples of interatomic interactions include hydrogen bonds and Van der Waals forces.

We performed analyses centered around small-worldliness as well as degree and closeness centrality distribution comparisons between both PCNs and RIGs. We analyze only the RIGs to confirm the observations seen in the prior Amitai et al. experiment [21]. We analyze the functional residues based on their closeness centrality and residue solvent accessibility (RSA) values. Functional residues are defined as binding sites for AcrIF1 and AcrIIA1 [22, 23]. For AcrVA1, functional residues are identified through loss-of-function mutagenesis studies, though their specific mechanistic roles are not yet known [24].

### 0.1 Measures of small worldliness are maintained between PCNs and RIGs

Small-world networks balance two properties - high clustering coefficient (implying dense local connections) and short average path length (implying efficient global connectivity). PCNs are commonly referred to as small-world networks [25, 26], but the same cannot be said for RIGs; we confirm this here.

In Table 1, we show the small worldliness values (sigma *σ*, normalized omega *ω_n_*, and small-world index/SWI) calculated for both PCN and the RIG for each Acr protein via a randomization test.

**Table 1.** Small worldliness measures (sigma, normalized omega, and small-world index/SWI) calculated for AcrIF1, AcrIIA1, and AcrVIA1.

| Small Worldliness Measures |  |  |  |  |
| --- | --- | --- | --- | --- |
| Protein | Number of nodes | Sigma | Normalized Omega | Small-world index |
| AcrIF1 (PCN) | 78 | 3.6014 | 0.8123 | 0.6884 |
| AcrIF1 (RIG) | 78 | 3.3740 | 0.9556 | 0.5990 |
| AcrIIA1 (PCN) | 292 | 9.2384 | 0.6106 | 0.6992 |
| AcrIIA1 (RIG) | 292 | 8.0936 | 0.7715 | 0.6272 |
| AcrVIA1 (in complex) (PCN) | 1260 | 7.1922 | 0.5891 | 0.6694 |
| AcrVIA1 (in complex) (RIG) | 1312 | 28.0531 | 0.7207 | 0.6649 |

All networks demonstrate small-world structure, with *σ* values ranging from approximately 3 to 28 across both network types, well above the threshold of *σ >* 1. It is important to note that the *σ* value is heavily dependent on the number of nodes (*n*) and edges (*k*), and has no mathematical maximum. That said, provided that AcrIF1 is the smallest structure, it has the smallest *σ* values, while AcrVIA1 has the largest *σ* values. When comparing the PCN and RIG values, we see that the PCN has larger *σ* values for AcrIF1 and AcrIIA1, but not for AcrVIA1.

*ω_n_* values are normalized following Neal’s method, where values closer to 1 indicate networks that approach the mathematical ideal of small-worldliness, with *ω_n_*= 1 representing perfect balance between clustering and path length [27]. We see that the both networks for AcrIF1 have the highest values, while this goes down for AcrIIA1 and AcrVIA1. We also see that the RIG network always has a higher *ω_n_*value than the PCN.

Finally, the SWI values range from 0 to 1, where 1 implies the graph is exhibiting both small-world characteristics (see Methods for more clarification). We see that with these values, these is overall less variation among the networks with the values being between 0.59 to 0.69. For AcrIF1 and AcrIIA1, we see that the RIG architectures have slightly lower SWI values than their PCN counterpart, but the difference between these values seemingly decreases as the size of the network increases (difference of 0.0894 between AcrIF1 PCN and RIG, difference of 0.072 between AcrIIA1 PCN and RIG, and 0.0045 between AcrVIA1 PCN and RIG).

All three indices demonstrate that PCNs and RIGs exhibit small-world properties. However, RIGs show consistently stronger small-worldliness across *ω_n_*values (ranging from 0.72 to 0.95). For the *σ* values, there is less consistency with the RIG showing noticeably stronger small-worldliness only for the largest protein graph (AcrVIA1). The difference between PCN and RIG SWI values decreases with increasing network size, raising questions about how graph construction methodology influences network topology.

### 0.2 Nodes representing the functional residues within PCNs and RIGs tend to have degrees closer to the mean degree

We created count distribution graphs (Figs 2, 3, and 4) to visualize what the average degree is for a residue within each protein’s PCN and RIG. The degree of a node is the number of connections that a node has to other nodes. In context of the PCN, a high degree node is a residue’s alpha carbon that has many other alpha carbons within 7 angstroms. In context of the RIG, a high degree node is a residue that has many interatomic contacts with other residues.

**Fig 2.**
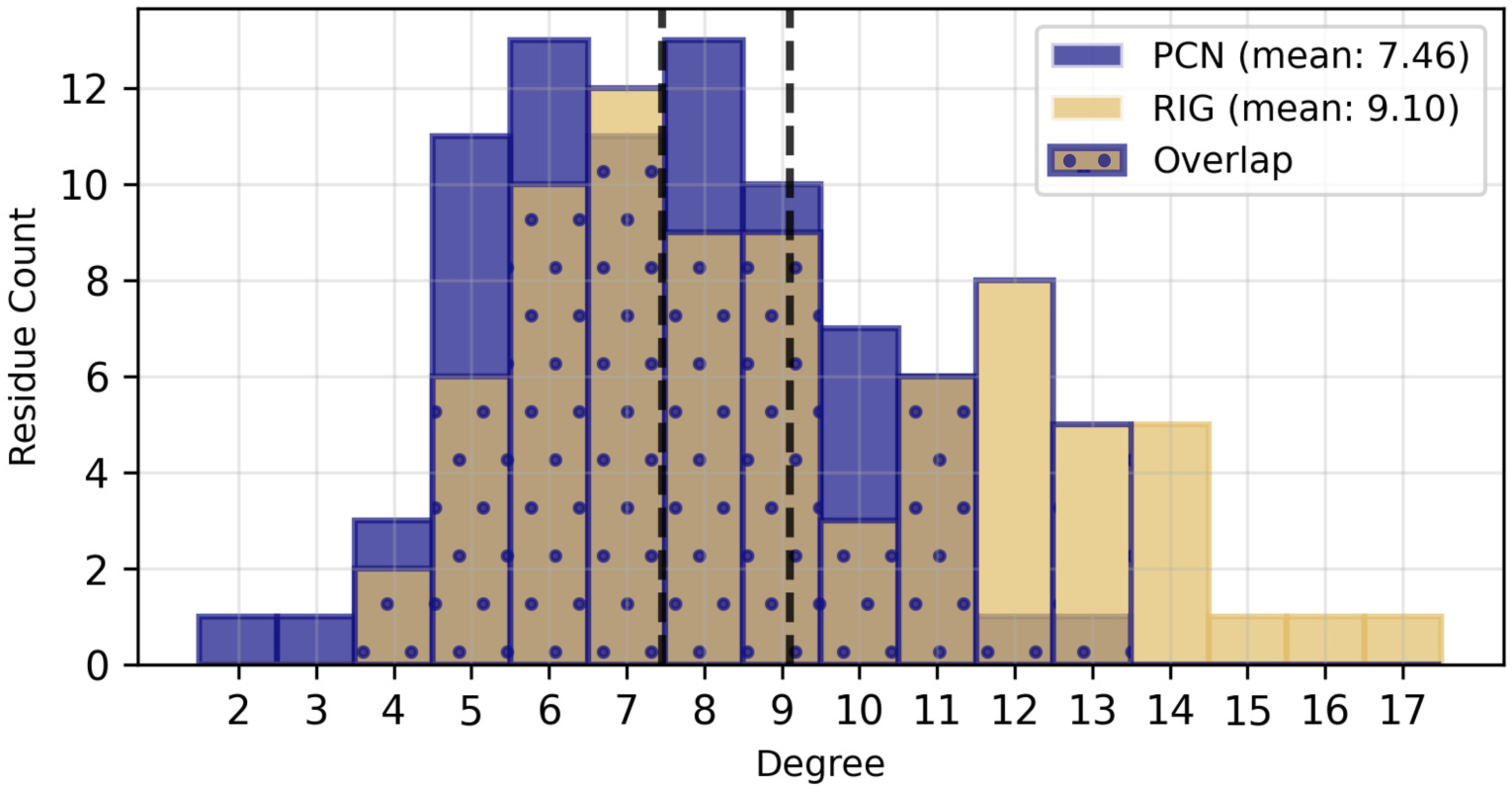
Comparison of degree distribution within protein contact network and residue interaction graphs of protein AcrIF1. Distribution of node degree for the protein contact network and residue interaction graph for protein AcrIF1, including the mean degree values for each network representation. The PCN has a lower mean degree of 7.46 compared to the mean degree of 9.10 for the RIG (mean values indicated by black dashed lines).

We outline the degrees for each of the nodes that correspond to a functional residue below. We note that generally, within a PCN, the degree of the functional residues are lower than within the RIG.

- AcrIF1
  – Tyr6 (PCN degree: 8, RIG degree: 7)
  – Tyr20 (PCN degree: 10, RIG degree: 11)
  – Glu31 (PCN degree: 10, RIG degree: 10)
- AcrIIA1
  – Phe115 (PCN degree: 6, RIG degree: 12)
- AcrVA1 (in complex)
  – Tyr39C (PCN degree: 4, RIG degree: 6)
  – Ser40C (PCN degree: 6, RIG degree: 9)
  – Asn43C (PCN degree: 7, RIG degree: 11)
  – Ser93C (PCN degree: 9, RIG degree: 11)
  – Gln96C (PCN degree: 8, RIG degree: 13)

In reference to the degrees of the functional residues outlined above, in Figs 2, 3, and 4, we show the count distribution of the degree of the residues, along with the average degree for both PCNs and RIGs. We can see in Fig 2 that for AcrIF1, the degree of the functional residues is only slightly above that of the mean for both the PCN and RIG. For AcrIIA1 in Fig 3, only the RIG degree of the single residue of interest is above the mean. And finally for AcrVA1 in Fig 4, three out of five total functional residues (Tyr39C, Ser40C, Asn43C) of interest are below the PCN degree mean, while two out of five functional residues are below the RIG degree mean.

**Fig 3.**
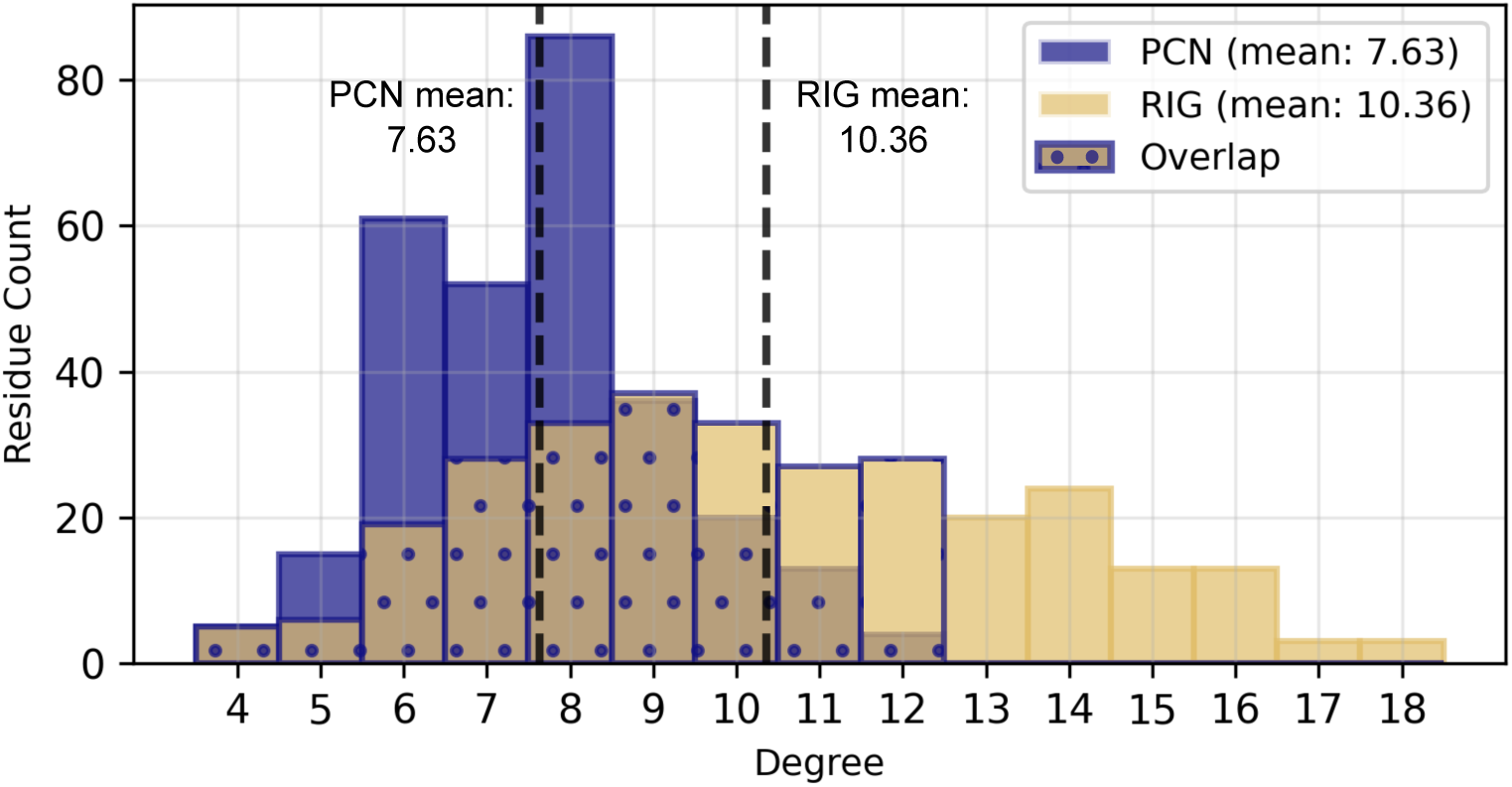
Comparison of degree distribution within protein contact network and residue interaction graphs of protein AcrIIA1. Distribution of node degree for the protein contact network and residue interaction graph for protein AcrIIA1, including the mean degree values for each network representation. The PCN has a lower mean degree of 7.63 compared to the mean degree of 10.36 for the RIG (mean values indicated by black dashed lines).

**Fig 4.**
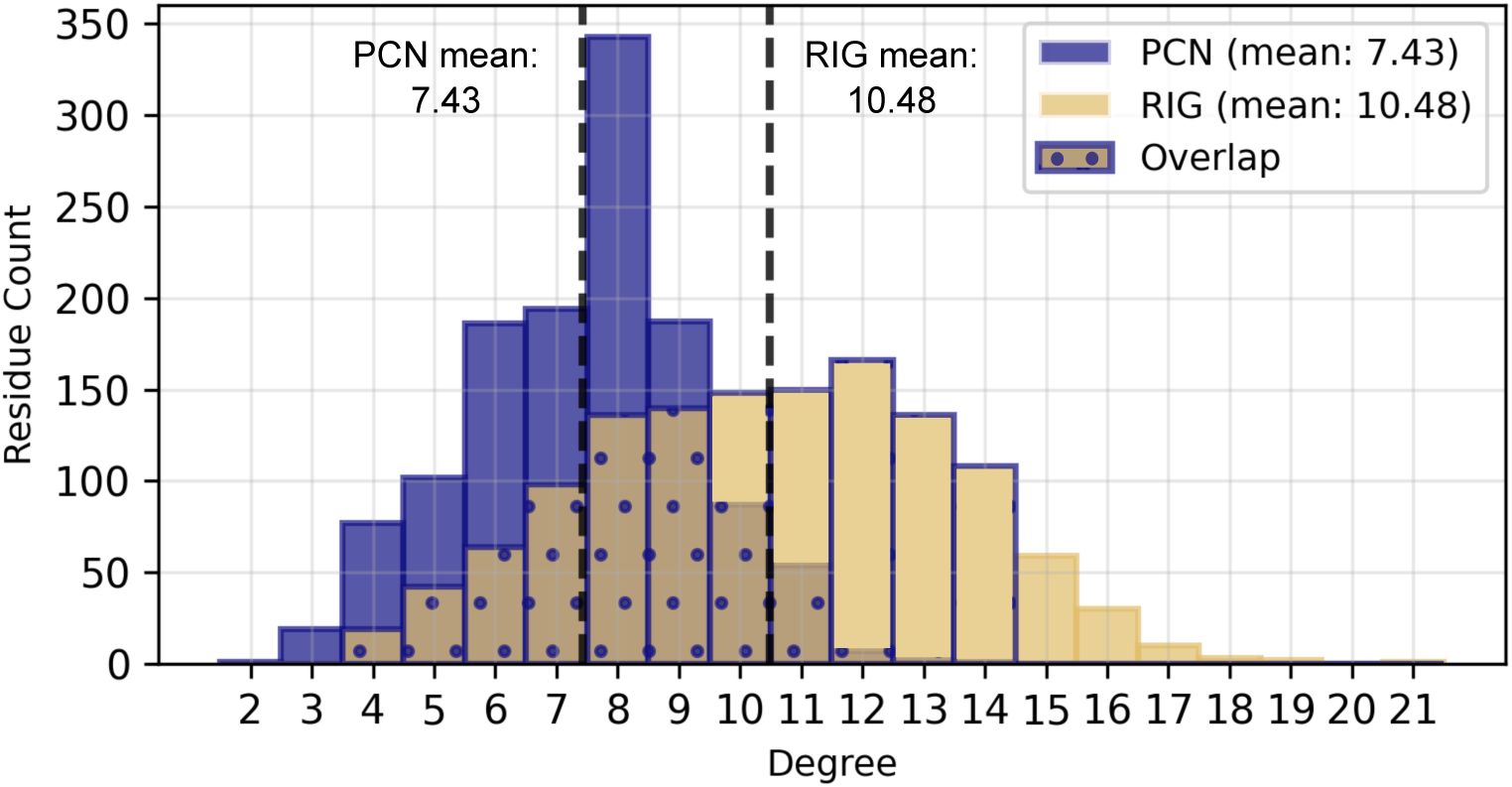
Comparison of degree distribution within protein contact network and residue interaction graphs of protein AcrVIA1 (in complex). Distribution of node degree for the protein contact network and residue interaction graph for protein AcrVIA1, including the mean degree values for each network representation. The PCN has a lower mean degree of 7.43 compared to the mean degree of 10.48 for the RIG (mean values indicated by black dashed lines).

Generally, it can be seen that the mean node degree is lower for all of the PCNs compared to the RIGs of the same protein. Interestingly, the mean degree for the PCNs are generally around 7 for all three proteins, and around 9-10 for the RIGs for all three proteins - this is regardless the total number of nodes within that protein’s RIG (see Table 1, Column 2). RIGs generally have more edges in comparison to PCNs as they include sidechain interactions. An amino acid can have various lengths of sidechains and these lengths are what contribute to why it is possible to have “distant” residues within a sequence that ultimately interact with each other within a protein. The architecture of a PCN is limited to establishing an edge only within a certain distance threshold of some individual and particular atom of interest (such as the alpha-carbon) - therefore, not accounting for these longer distance interactions. As RIGs have more edges, their nodes also tend to have higher degrees.

### 0.3 Nodes within PCNs and RIGs tend to have closeness centralities either less than or closer to the mean closeness centrality

We created count distribution graphs (Figs 5, 6, 7) to visualize what the average closeness centrality is for a residue within each protein’s PCN and RIG. Closeness centrality is a distance measure, calculated as the reciprocal of the sum of the length of the shortest path between the node and any other nodes within that connected graph. A larger value for closeness centrality indicates a more central node which would be able to interact with more nodes compared to a less central node. In context of the PCN, a high closeness centrality node is a residue’s alpha carbon that is spatially positioned such that it can reach many other alpha carbons through fewer intermediate contacts.

**Fig 5.**
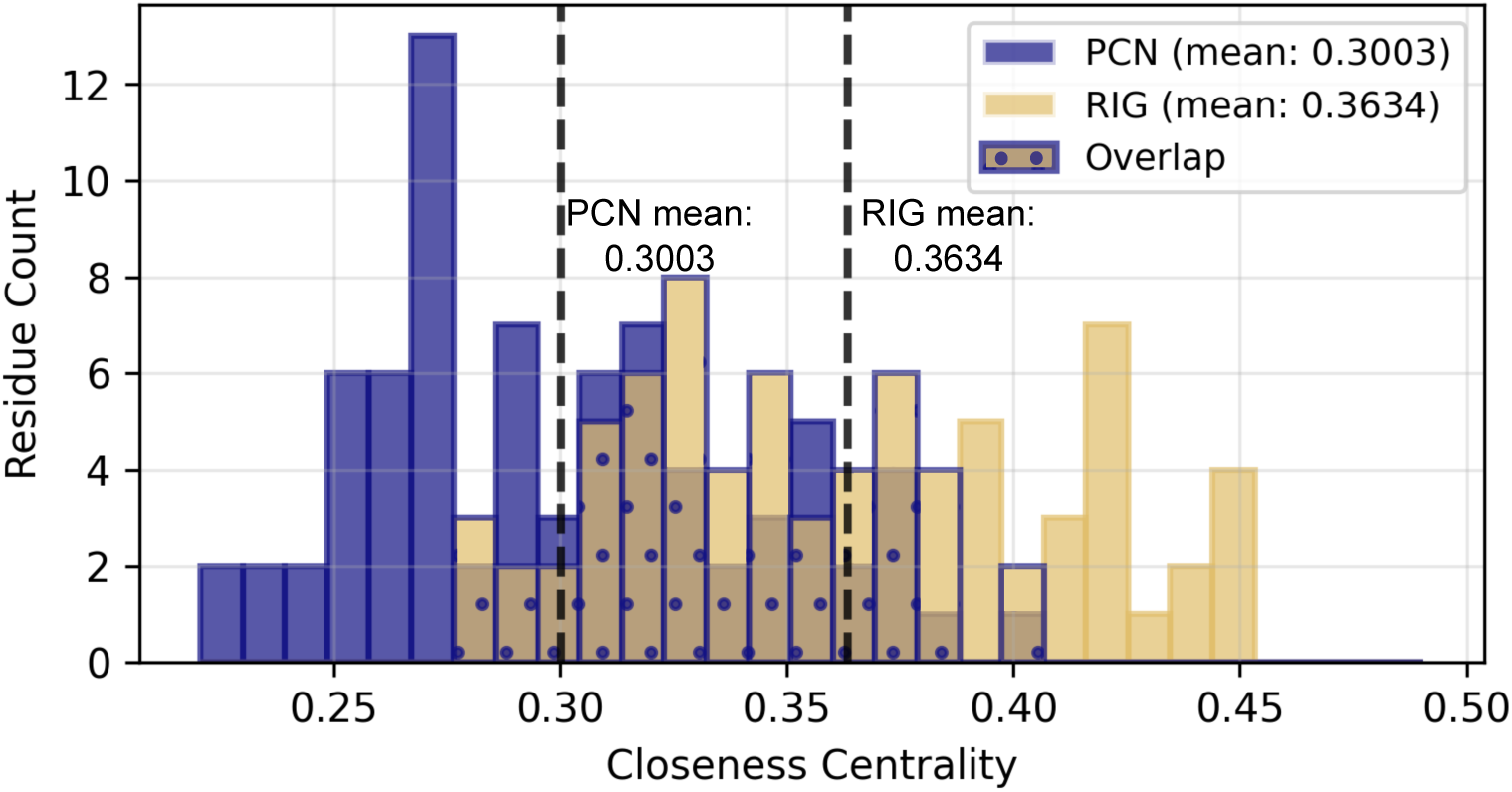
Comparison of closeness centrality distribution within protein contact network and residue interaction graphs of protein AcrIF1. Distribution of node closeness centrality for the protein contact network and residue interaction graph for protein AcrIF1, including the mean degree values for each network representation indicated by black dashed lines. The PCN has a lower mean closeness centrality (0.3003) compared to the corresponding RIG value (0.3634).

In context of the RIG, a high closeness centrality node is a residue that is well-positioned within the web of interatomic interactions, requiring fewer intermediary residues to reach any other residue in the network.

- AcrIF1
  – Tyr6 (PCN closeness centrality: 0.3333, RIG closeness centrality: 0.3719)
  – Tyr20 (PCN closeness centrality: 0.3333, RIG closeness centrality: 0.3869)
  – Glu31 (PCN closeness centrality: 0.3581, RIG closeness centrality: 0.4162)
- AcrIIA1
  – Phe115 (PCN closeness centrality: 0.1082, RIG closeness centrality: 0.1707)
- AcrVA1 (in complex)
  – Tyr39C (PCN closeness centrality: 0.0602, RIG closeness centrality: 0.1242)
  – Ser40C (PCN closeness centrality: 0.0602, RIG closeness centrality: 0.1251)
  – Asn43C (PCN closeness centrality: 0.0594, RIG closeness centrality: 0.1232)
  – Ser93C (PCN closeness centrality: 0.0753, RIG closeness centrality: 0.1381)
  – Gln96C (PCN closeness centrality: 0.0720, RIG closeness centrality: 0.1374)

We show the count distribution of the closeness centrality of the residues, along with the average closeness centrality for both the PCN and RIG. We can see in Fig 5 that for AcrIF1, the closeness centrality of the functional residues is only slightly above that of the mean for both the PCN and RIG. For AcrIIA1 in Fig 6, neither the PCN nor the RIG closeness centrality of the single residue of interest is above the mean. And finally for AcrVA1 in Fig 7, three residues out of the total five (Tyr39C, Ser40C, Asn43C) are below the mean for the PCN, while all of the residues are either slightly below or above the RIG mean.

**Fig 6.**
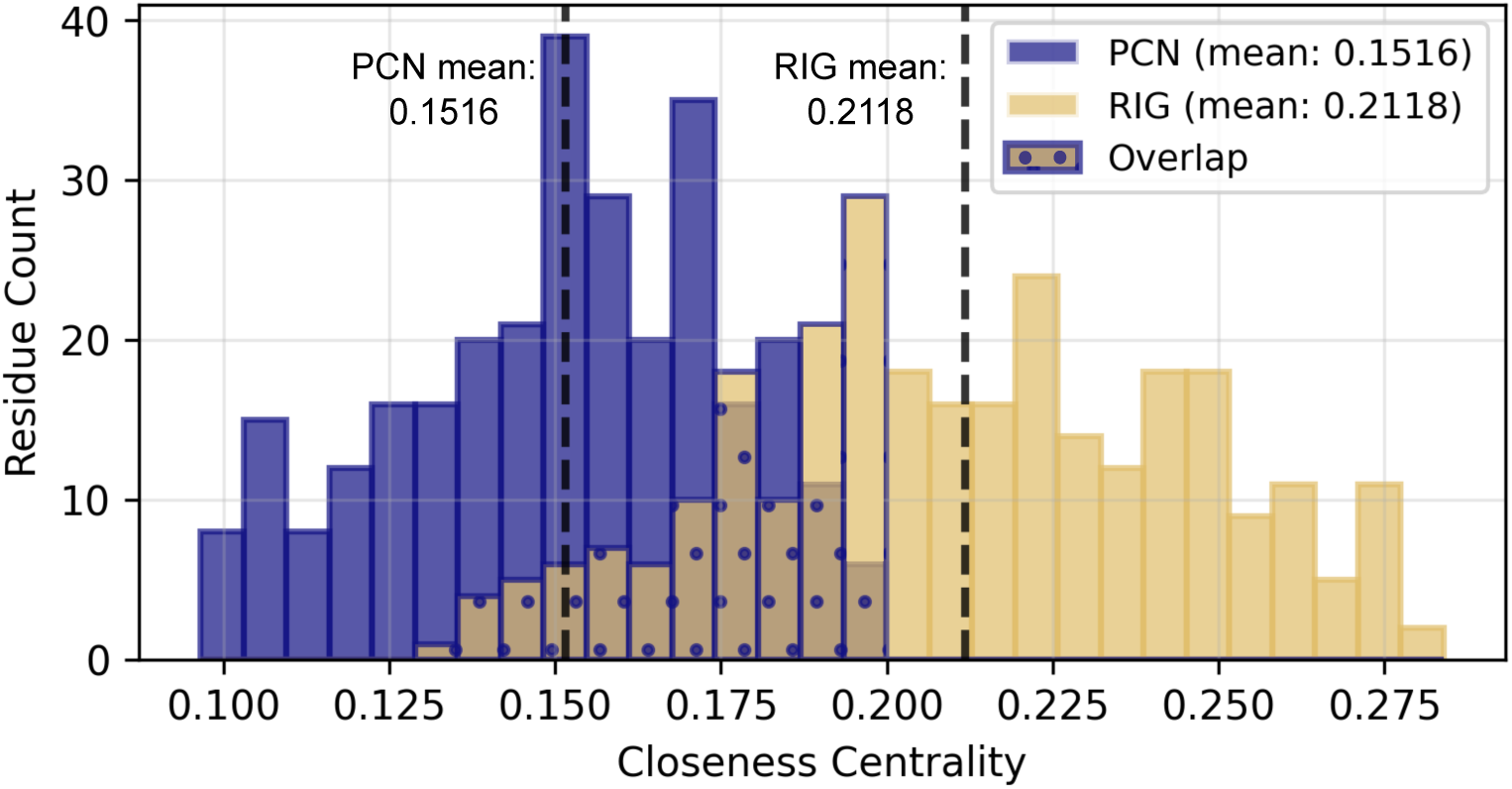
Comparison of closeness centrality distribution within protein contact network and residue interaction graphs of protein AcrIIA1. Distribution of node closeness centrality for the protein contact network and residue interaction graph for protein AcrIIA1, including the mean degree values for each network representation indicated by black dashed lines. The PCN has a lower mean closeness centrality (0.1516) compared to the corresponding RIG value (0.2118).

**Fig 7.**
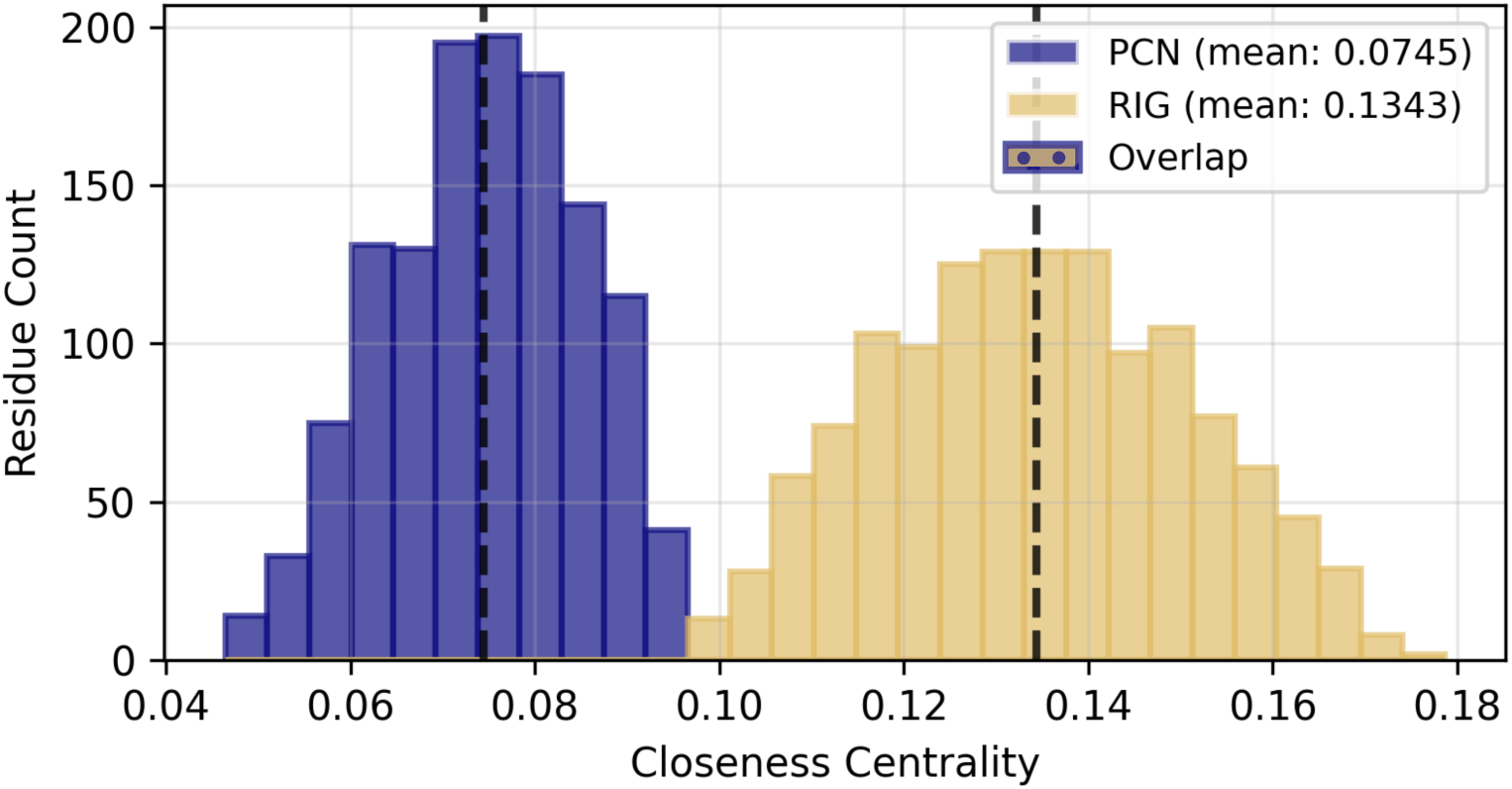
Comparison of closeness centrality distribution within protein contact network and residue interaction graphs of protein AcrVA1 (in complex). Distribution of node closeness centrality for the protein contact network and residue interaction graph for protein AcrVA1, including the mean degree values for each network representation indicated by black dashed lines. The PCN has a lower mean closeness centrality (0.0745) compared to the corresponding RIG value (0.1343). We observe that there is no overlap of values between the PCN and RIG, unlike the other proteins.

There is noticeably no overlap of closeness centrality values for AcrVA1 unlike the other graphs, though this is possibly related to the differences between the architectures of the PCN and RIG. Additionally, there is a decreasing trend in the closeness centrality values as the PCNs and RIGs get larger (i.e. more nodes and edges), with the RIGs generally always having a higher mean closeness centrality value than the PCN. The RIG, with more nodes and more edges, may include more connections that skew the closeness centrality measures to be higher, reflecting generally better connected nodes with shorter length paths to other nodes.

### 0.4 Functional residues for AcrIF1, AcrIIA1, and AcrVA1 in RIGs show high variability in closeness centrality and RSA values

Originally Amitai et al. demonstrated that high closeness values can be associated with functional residues [21]. Additionally, they demonstrated that these high closeness residues also tend to have low residue solvent accessibility (RSA) values ranging from 20% to 40%, meaning these residues are less exposed [21].

We utilize the RIGs created for AcrIF1, AcrIIA1, and AcrVA1 and determine standardized values based off of amino acids found in 2688 PDB files of proteins that were all transformed into RIGs. We curated this set of proteins of have similarly sized RIGs to that of Acr proteins - that is, Acr proteins generally have about 50 to 200 or 300 amino acids [28, 29]. We discuss the method for this and which was used in Amitai et al. in Section 0.8 of the Methods. We map all residues and their corresponding standardized degree and closeness centrality in Figs. 8, 9, and 10.

**Fig 8.**
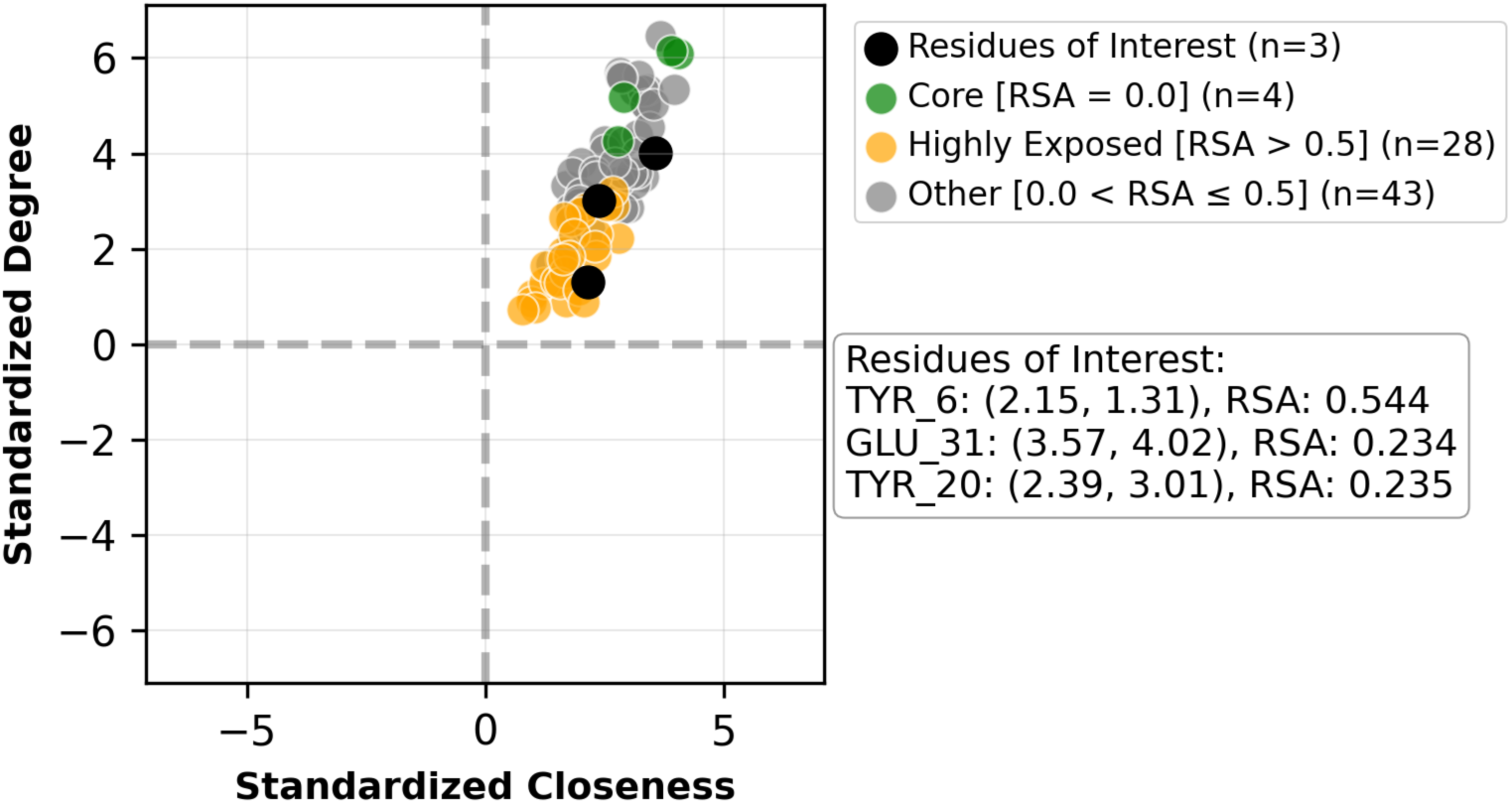
Standardized degree vs. standardized closeness centrality for residues based on the protein AcrIF1 RIG. Values are shown for all 78 nodes within the AcrIF1 RIG. Highly exposed residues (RSA *>* 0.5) are in yellow, core residues (RSA = 0.0) are in green, and all other residues are in grey. See Methods for how degree and closeness centrality values are standardized. We can see that out of the three residues, only Tyrosine 6 (TYR6) has an RSA over 0.5, which would be considered highly exposed. The residues in AcrIF1 tend to skew higher than the average in degree and closeness centrality.

**Fig 9.**
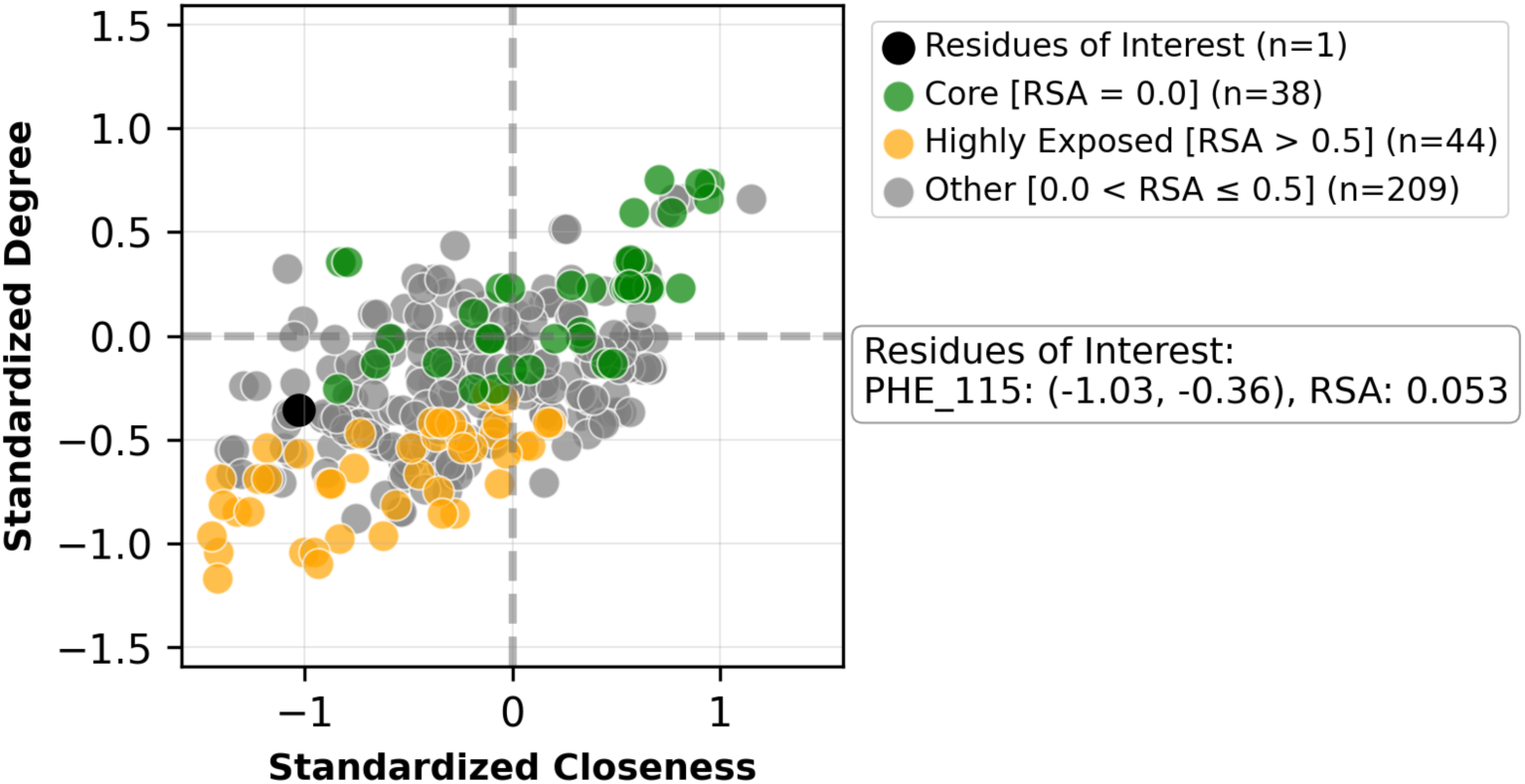
Standardized degree vs. standardized closeness centrality for residues based on the protein AcrIIA1 RIG. Values are shown for all 292 nodes within the AcrIIA1 RIG. Highly exposed residues (RSA *>* 0.5) are in yellow, core residues (RSA = 0.0) are in green, and all other residues are in grey. See Methods for how degree and closeness centrality values are standardized. There is only one listed residue of interest, Phenylalanine 115, and its RSA is around 5%, indicating it is less exposed. There is more spread in terms of standardized degree and closeness centrality values among the residues compared to residues in AcrIF1 and AcrVA1, with many highly exposed residues often having lower degree or closeness centrality.

**Fig 10.**
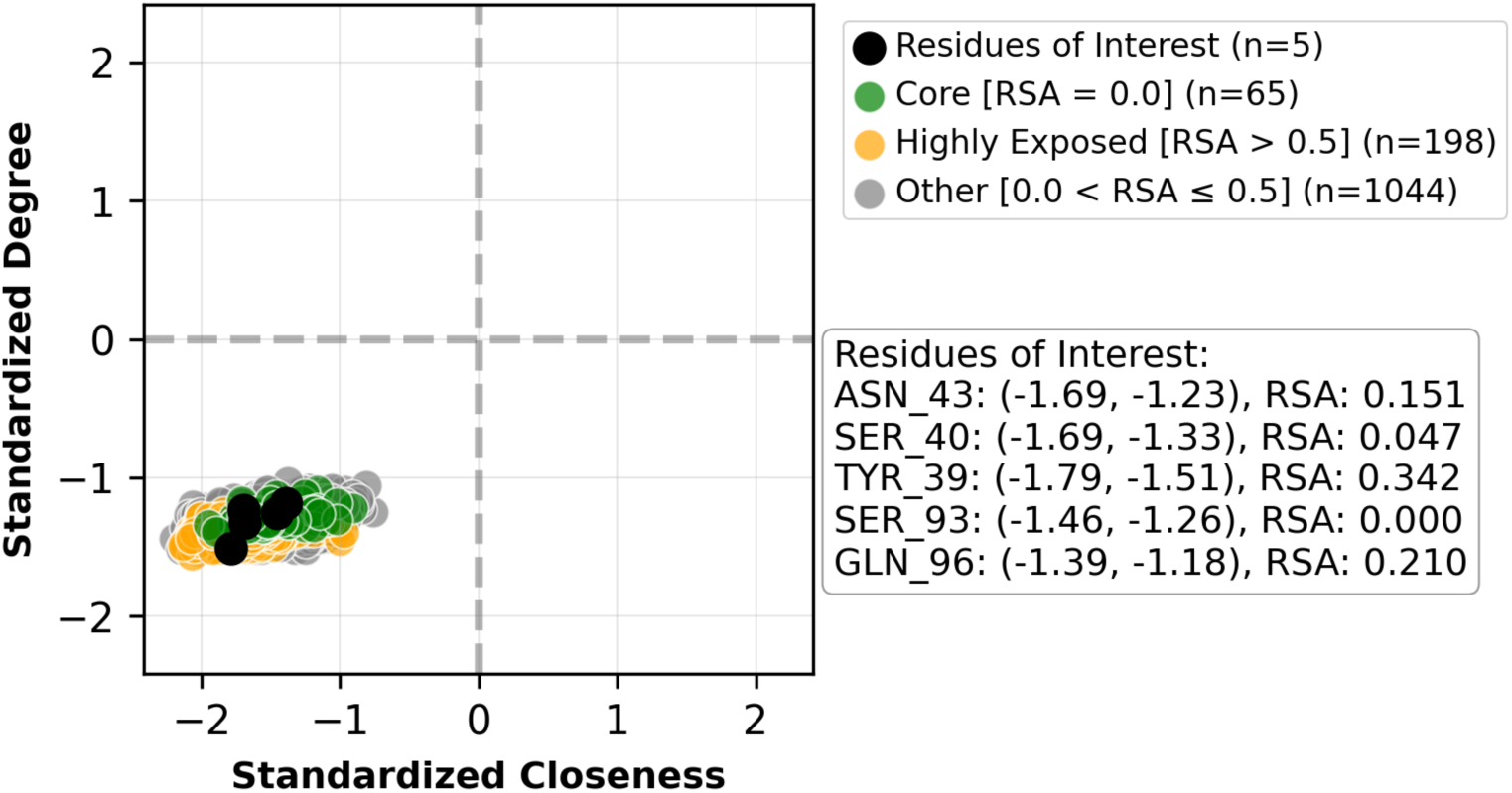
Standardized degree vs. standardized closeness centrality for residues based on the protein AcrVA1 (in complex) RIG. Values are shown for all 1260 nodes within the AcrVA1. Highly exposed residues (RSA *>* 0.5) are in yellow, core residues (RSA = 0.0) are in green, and all other residues are in grey. See Methods for how degree and closeness centrality values are standardized. There are five functional residues (Asparagine 43, Serine 40, Tyrosine 39, Serine 93, and Glutamine 96), but none of these are considered to be highly exposed residues based on RSA values. All of the residues have lower than average degree and closeness centrality values within the RIG for AcrVA1.

There is overlap between these results and what was determined in Amitai et al. of the highly exposed residues (seen in yellow) generally have the lowest values for standardized degree and for standardized closeness, while core residues generally have some of the higher values. However, the conclusions that showed that many functional residues (i.e. residues that contribute to the function of the protein) can generally have high closeness centrality does not always hold true for these three proteins, regardless of the PCN or RIG being used as the representation.

## Discussion

Just as network science has been used to represent complex systems on the scale of public transport systems within cities to determine how accessible some routes may be over others [30], it has and will continue to be used on a smaller scale to model proteins and as a basis for other *in silico* methods. Our work further refines and clarifies the use cases of this approach to smaller proteins through a case study of three niche viral proteins, as well as what implications should be accounted for.

We constructed both protein contact networks (PCNs) and residue interation graphs (RIGs) for three anti-CRISPR (Acr) proteins - Acr30-35 or AcrIF1 (PDB ID: 2LW5, solution NMR), AcrIIA1 (PDB ID: 5Y6A, x-ray diffraction), and AcrVIA1 (PDB ID: 6VRB, electron microscopy). We must state the following limitation: the method through which the structure is obtained may affect the construction of the PCN or RIG. Generally, x-ray diffraction provides greater resolutions at the expense of the protein being crystallized or stationary in some material. By keeping the protein crystallized, it may not reflect the natural conformation of the protein due to the preparation needed to create the image. In contrast, electron microscopy (EM) is commonly used to capture larger complexes of proteins at the expense of resolution. Additionally, scientists have used EM imaging for studies in dynamics, as one can see the protein in its soluble form, and thus, its actual conformations in solution. Both x-ray diffraction and EM are dependent on Fourier optics and reconstruction control. Finally, solution nuclear magnetic resonance (solution NMR) is another means of obtaining protein structures by measuring how nuclei respond to radio frequency pulses. Resolution quality determines how well the atoms can be distinguished – the poorer the quality, the less atoms can be distinguished meaning that more of the atomic structure must be hypothesized. This can likely affect any algorithm used to then build a graph reliant on positional data from the PDB file.

Anti-CRISPR (Acr) proteins, as they are so newly discovered, are limited in the amount of structural data available, especially in terms of single, x-ray diffraction structures which were originally used in Amitai et al. [21]. Further research studies should select for an individual protein which has structural data for all of three of the above described methods and compare the differences in PCN and RIG construction. Those studies could provide more detailed explanation to these hypothesized differences between these methods commonly used within structural biology.

We confirmed the small worldliness of each network based on updated measures referenced by Neal [27], as studies of protein networks often cite small worldliness as a general property without explicitly quantifying or measuring this topology. The variation in small-worldliness metrics between network types seen in Table 1 warrants examination of their underlying structural differences. RIG architectures consistently exhibit higher edge density than PCN architectures. This difference stems from the fundamentally different construction methods: PCN distance-based graphs use a binary cutoff applied to a single point per residue (typically the alpha carbon), creating an edge whenever two alpha carbons fall within the specified distance threshold. In contrast, RIG contact-based graphs are more inclusive—each residue contains multiple atoms (backbone and side chains), providing numerous potential contact points between residues. This multi-point contact criterion captures more interactions than the single-point distance threshold, resulting in denser RIG networks.

Small worldliness has been linked to diverse phenomena, from the technical and commercial success of open-source software [31] to information transfer within the brain [32]. It has also been analyzed specifically within proteins, revealing links between residues and protein dynamics [25]. This broad relevance motivates further investigation into whether small worldliness can provide additional insights into protein structure networks of small viral proteins through the spatial positioning of residues or the interactions among residues themselves.

We applied an altered portion of a methodology previously utilized in larger proteins [21]. This methodology predicts that functional residues should exhibit high degree and closeness centralities, as these residues are expected to“integrate and transmit” information throughout the protein [21]. However, even with a curated set of proteins of similar sizes to anti-CRISPR proteins, we did not observe this pattern.

Amitai et al. discussed lack of accuracy in their techniques for their smallest proteins (less than 120 residues). The paper originally used a Jackknife resampling technique to determine optimal thresholds for closeness centrality and RSA that could characterize important residues. Jackknife resampling estimates the variability of a statistic by iteratively excluding each data point one at a time and recalculating the statistic on the remaining data. In the context of protein networks, this means recalculating threshold values after sequentially removing each protein from the dataset, which can be problematic when network sizes are highly variable. Smaller protein networks have many nodes with similar values that may skew above or below the average (Figs. 8, 9). Thus, if the Jackknife technique results in a sufficiently high threshold, none of the residues of this protein would be considered functional.

Since the Jackknife technique could produce thresholds that are either too high or too low, it would be beneficial to evaluate the data used in the standardization. The Amitai et al. experiment focused on three proteins of which one is a bacterial protein (subtilsin), and two are eukaryotic proteins (ERK2 MAP kinase and glycogen phosphorylase). It used 42 protein chains from various sources (ranging from *Escherichia coli* to the *Rattus norvegicus* or the brown rat) to provide standardized values for the residues. Additional research studies could explore to what extent these standardized values changes if residues are sourced from primarily eukaryotic, prokaryotic, or viral proteins or a more narrow subset of classifications.

The challenges increase when these graph representations are used as input for deep learning models. Deep learning methods for protein function prediction, such as DeepFRI, rely on graph representations of protein structure as primary input. DeepFRI combines a graph convolutional neural network (GCN) with large language model (LLM) features, using PCNs as the structural input [20]. However, GCNs exhibit a known bias toward high-degree nodes in node classification tasks, resulting in higher misclassification rates for low-degree nodes [33, 34]. This presents a potential problem: our results show that functional residues within Acr proteins often have low or average degrees in both PCN and RIG representations.

This mismatch between model bias and biological reality may be exacerbated by the graph construction method itself. PCN distance thresholds that use only the alpha carbon of a residue can miss biologically relevant interactions between residues that are distant in sequence but spatially proximate due to varying side chain lengths. These “distant” sequence interactions contribute to protein function but may be underrepresented in PCN architectures, compounding the challenge of identifying low-degree functional residues.

Two potential solutions exist: either adapt the data representation to better suit existing models (rethink PCN and RIG architectures and explore different distance thresholds), or adapt the model architectures to better handle the data characteristics (such as using GNN variants with reduced degree bias). For small bacterial and viral proteins like Acrs, exploring alternative GNN architectures or enhanced graph representations may improve functional residue prediction. Further experimentation building upon existing models like DeepFRI, with careful consideration of the underlying network structures used as input, could expand their applicability to these challenging protein classes.

In summary, our analysis of anti-CRISPR (Acr) proteins reveals important considerations for applying network-based methodologies to small proteins. While some techniques have proven valuable for analyzing larger proteins, the structural and topological characteristics of smaller proteins (particularly the tendency of the nodes of functional residues to exhibit low degrees) present distinct challenges for graph-based computational methods. The bias of GCN architectures toward high-degree nodes, combined with the rigid distance thresholds inherent to PCN construction, suggests that current deep learning approaches may systematically underperform on small viral and bacterial proteins. These findings prompt the need for methodological adaptations such as using degree-aware or alternative graph neural network architectures, establishing protein-type-specific standardization values, and accounting for the unique structural properties that emerge at smaller scales. As computational methods increasingly rely on structural networks as input, understanding these scale-dependent limitations becomes essential for accurate functional prediction across the full spectrum of protein sizes.

## Methods

We outline our process as the following: the selection of the proteins of interest, the construction of the protein contact networks (PCNs) and residue interaction graphs (RIGs), and the analysis of the PCNs and RIGs utilizing techniques from the Amitai et al. experiment [21]. We also provide information regarding the idea of “small-world” networks, which is the designation often provided for graphs utilizing protein structural data.

### 0.5 Selection of model Acr proteins

We have narrowed our focus to three experimentally validated Acrs as they have known and experimentally validated essential residues: AcrIF1 (also known as Acr30-35), PDB ID: 2LW5), AcrIIA1 (PDB ID: 5Y6A), and AcrVIA1 (PDB ID: 6VRB, in complex).

The test sequences contain 78 (AcrIF1), 302 (AcrIIA1), and 229 (AcrVIA1) residues. Additional information about these Acr proteins can be found in the Bondy-Denomy database [35].

In reference to prior work, these network analyses applied to monomeric and dimeric structures created via X-ray diffraction which generally have a resolution of about 2 angstroms [36]. We expand on this by exploring both monomeric (AcrIF1), dimeric (AcrIIA1), and polymeric (AcrVIA1) structures, as well as utilizing PDB data acquired through EM and solution NMR.

### 0.6 Construction of protein structure networks and residue interaction graphs

We use two different network representations for proteins – the protein contact network (PCN) and the residue interaction graph (RIGs). Bhattacharyya et al. [16] contains a chart that details different naming conventions, definitions, and applications of network representations for proteins. They describe a residue interaction graph (RIG) as a subtype of a protein structure network (PSN); however, to better clarify to the reader which architecture refers to which in this paper, and to maintain current used nomenclature, we describe each as follows:

- Protein contact network (PCN) architecture denotes a node as a residue’s alpha or beta carbon, and the edges exist if two nodes are found to be within some distance threshold of each other. We are using the alpha carbons and a distance of 7 angstroms (a standard distance used for alpha carbon interactions, but can otherwise, differ).
- Residue interaction graph (RIG) architecture denotes a node as a residue, and the edges exist if there are interactions between the nodes. Interactions include backbone peptide bonds and non-covalent bonds (i.e. hydrogen and hydrophobic interactions), often determined using third-party software.

To create the PCNs, we utilize Biopython to parse the information from the PDB files as well as creating our own functions using the Networkx libraries. Those associated functions can be found in the Github. To create the RIGs, we utilize PDBe-Arpeggio [37], which calculates interatomic contacts based on rules defined in CREDO, a publicly available database of protein-ligand interactions [38].

It is important to clarify that regarding the structures of the PCN and RIG for AcrVA1, these graphs are that of a protein complex (the Acr protein, a Cas13 protein, and RNA). We omitted the RNA from being included as a node within the PCN. However, the RNA was not omitted in terms of the RIG construction as Arpeggio will consider RNA when determining interactions, as described above. This means that the RIG will inherently have more nodes and edges compared to the PCN for this structure in particular. Likewise, we did not omit the Cas13. It is not practical to omit the Cas13 and RNA since the AcrVA1 protein structure is conformed with it and not in a solitary, natural state. Within supplemental files for this complex, AcrVA1 is associated with chain C, and associated residues include a C to clarify this.

AcrIF1 and AcrIIA1, in contrast, are within a solitary, natural state. Therefore, these networks contain the same amount of nodes between the PCN and the RIG, but not necessarily the same amount of edges. In this experiment, the RIGs always have more edges than the corresponding PCN.

### 0.7 Calculating residue solvent accessibility

Residue solvent accessibility (RSA) is a measure of the exposure of that residue within the 3D structure, often used to help describe a protein’s biophysical or evolutionary properties [39]. To calculate RSA, first we calculate the solvent-accessible surface area (SASA) using Biopython’s implementation of the Shrake and Rupley algorithm [40]. The formula for the RSA calculation can then be found

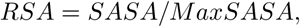

where MaxSASA is the maximum possible solvent accessible surface area for that residue. We utilized the empirical values for MaxSASA found in Tien et al. (2013) [39]. The RSA measure is read as a percentage, where a value of 0.0 would indicate a residue is not exposed or otherwise, core. A RSA value greater than 0.5 (or 50%) means that the residue is considered to be highly exposed.

### 0.8 Revisiting the Amitai et al. experiment

This experiment uses a portion of the methodology used by Amitai et al. [21] where they demonstrated that for larger proteins (subtilsin, ERK2 MAP kinase, and glycogen phosphorylase), residue interaction graphs (RIGs) can be used to predict functional residues. They explored various network centrality metrics such as degree centrality (the amount of connections nodes have within the network) and closeness centrality (the distance of a node to other nodes within the network), as well as utilizing residue solvent accessibility (RSA). Following the section denoted “Transforming protein structures into residue interaction graphs”, we sourced proteins with the following properties:

- less than 30% sequence identity between the proteins
- resolutions less than 3 angstroms
- no chain breaks
- no mutants
- no membrane proteins

We also added a sequence length limit of 40-500 residues (as one shared characteristic among Acr proteins is their small size - about 50 to 200 or 300 amino acids [28, 29]), and an R-value of 0.25 or less. As explained by Protein Data Bank (PDB), R-value is a measure of the quality of the atomic model. A random set of atoms would have an R-value of about 0.63, while a perfect fit would have a value of 0. The typical values for structure would be about 0.20.

The one exception made from the properties originally proposed by Amitai et al. above was the B-factor *<* 0.3 requirement. B-factor, also known as the Debye-Waller factor, describes the reduction of X-ray or neutron scattering caused by thermal motion and can be used to identify the flexibility of atoms, sidechains, or regions [41]. A lower B-factor within an atom generally indicates more order or otherwise, reliable positioning within x-ray crystallography data. That said, we looked at the validation reports available on the PDB entry website for each of the Acr proteins - only AcrIIA1 (PDB ID: 5Y6A) has a B-factor as it was gathered using x-ray diffraction. B-factor is not used with solution NMR and its definition is different in electron microscopy. These validation reports are publicly available on the PDB page for each respective protein, and can be searched using the PDB ID.

We sourced 2684 PDB files using the PISCES server [42]. More information about the proteins used (PDB ID, organism, etc.) is provided within the Github. The original Amitai et al. experiment used a different program (LPC CSU) to find all the interactions between residues [43]. As LPC CSU is available but unable to be used with these PDB files (possibly as it was originally made in 1999), we utilized PDBe-Arpeggio [37] to determine the non-covalent interactions as it is more recently made. Additionally, PDBe-Arpeggio also includes its own means of resolving issues that may come from atom flexibility (B-factor) when attempting to accurately determine interatomic contacts, which it discusses within its paper [37].

### 0.9 Confirmation of small-world property among Acr proteins

Amitai et al. begins with the assumption that their protein RIGs are small world networks. Given changes in quantification of small worldliness, and as the original experiment did not contain this information, we provide background details and explanations to what is meant by the term small-world network.

In 1998, Watts and Strogatz identified a network structure which is now associated with the term “small world” network [44]. They offered an example of a graph containing *n* vertices that are each connected to its *k* nearest neighbor via undirected edges. Choosing a random vertex and an edge as described above, they can reconnect an edge to a randomly chosen vertex with probability *p* (duplicate edges are not allowed). They continue to do this in a clockwise fashion around the graph for one lap.

Then, they look at the edges that connect the vertex to its second-nearest neighbor, third-nearest, and so on (for *k* /2 laps). For a *p* = 0, the graph is unchanged while for a *p* = 1, all the graph’s edges are rewired randomly. Small-world networks then are the intermediate graphs where 0 *< p <* 1.

There are various indicators used to measure small worldliness, though it is primarily quantified and computed from the observed network’s clustering coefficient (C) and the mean path length (L). Provided that the Amitai et al. experiments occurred in 2004, three main formulas have since come into use and the main difference is how they are normalized with respect to values from one or more reference graphs (random and/or lattice graph):

- Sigma (*σ*), or otherwise referenced as the small-world quotient (*Q*). The expression is as follows:

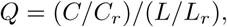

where *C_r_*and *L_r_*denote the clustering coefficient and mean path length of a single reference graph.
- Omega (*ω*), which rather than using a single reference graph, uses both a random and lattice reference graph. The expression is as follows:

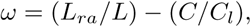

where *L_ra_* denotes the mean path length of the random reference graph and *C_l_* denotes the clustering coefficient of the lattice reference graph.
- Small-world index (SWI), which tries to account for variations in *C_ra_* rand *C_l_*, as well as *L_ra_* and *L_l_*. The expression is as follows:

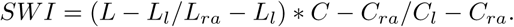

It looks for the maximization of both relationships.

They are characterized by the following properties:

- preserved local neighborhoods (in other words, tight connections exist between nodes which cluster).
- an average shortest distance that logarithmically increases with *n* (in other words, one can reach any node with few connections). Facebook demonstrated that among 1.59 billion people with active accounts on their social media platform, each person is connected to every other person by an average of three and a half people [45].

To get the values seen in Table 1, using NetworkX, we constructed 100 random and lattice reference graphs that contain the same amount of nodes and edges relative to the original PCN or RIG we are trying to analyze, and then used them to determine *C_r_*, *L_r_*, *C_ra_*, *L_ra_*, *C_l_*, and *L_l_* values. We then used the equations and definitions previously outlined in this section.

### 0.10 Disclosure of AI use

Claude Sonnet 3.7 and Claude Sonnet 4 were utilized throughout the methodology in the following ways: assistance with data organization (i.e., PDB files to CSV files with data), figure generation (matplotlib), for help with L^A^T_E_X formatting for tables or equations (i.e., Table 1), coding/commenting (i.e., creating functions to enable batch processing and increase efficiency of larger functions, creation of helper functions to utilize package functions on existing data, descriptions of functions), and general debugging purposes (i.e. package version issues in venv). Output of Claude Sonnet 3.7 and Claude Sonnet 4 were reviewed by the first author.

### 0.11 Data availability

All code is publicly accessible and commented at https://github.com/michramsa/AcrNetworks-clean. Majority of data is available at https://doi.org/10.5281/zenodo.17415956. As stated within the Github and Zenodo repositories, unavailable data is primarily due to size restrictions. This data can be generated by the code to the user’s local machine, but if access is needed to the exact files, contact the corresponding author and this data can be provided.

## Notes

### Competing Interest Statement

The authors have declared no competing interest.

